# Error-corrected Duplex Sequencing enables direct detection and quantification of mutations in human TK6 cells with remarkable inter-laboratory consistency

**DOI:** 10.1101/2023.02.22.529418

**Authors:** Eunnara Cho, Carol D. Swartz, Andrew Williams, Miriam Rivas, Leslie Recio, Kristine L. Witt, Elizabeth K. Schmidt, Jeffry Yaplee, Thomas H. Smith, Phu Van, Fang Yin Lo, Charles C. Valentine, Jesse J. Salk, Francesco Marchetti, Stephanie L. Smith-Roe, Carole L. Yauk

**Affiliations:** Environmental Health Science and Research Bureau, Health Canada, Ottawa, ON, Canada; Department of Biology, Carleton University, Ottawa, ON, Canada; Inotiv-RTP, Research Triangle Park, Durham, NC, USA; Scitovation, Research Triangle Park, Durham, NC; Division of Translational Toxicology, National Institute of Environmental Health Sciences, Research Triangle Park, Durham, NC, USA; TwinStrand Biosciences, Inc., Seattle, WA, USA; Department of Biology, University of Ottawa, Ottawa, ON, Canada

**Author notes:** Correspondence: Carole L. Yauk, Department of Biology, University of Ottawa, 269 Gendron Hall, 30 Marie-Curie Private, Ottawa, Ontario K1N 6N5, Canada., Correspondence: Stephanie L. Smith-Roe, Systems Toxicology Branch, Division of Translational Toxicology/NIEHS, PO Box 12233, MD-K17, Research Triangle Park, NC 27709. Equal contribution (Eunnara Cho, Carol D. Swartz).

**Keywords:** *N*-ethyl-*N*-nitrosourea, in vitro, error corrected next generation sequencing, mutagenesis, mutation spectrum

## Abstract

Error-corrected Duplex Sequencing (DuplexSeq) enables direct quantification of low-frequency mutations and offers tremendous potential for chemical mutagenicity assessment. We investigated the utility of DuplexSeq to quantify induced mutation frequency (MF) and spectrum in human lymphoblastoid TK6 cells exposed to a prototypical DNA alkylating agent, *N*-ethyl-*N*-nitrosourea (ENU). Furthermore, we explored appropriate experimental parameters for this application, and assessed inter-laboratory reproducibility. In two independent experiments in two laboratories, TK6 cells were exposed to ENU (25-200 µM) and DNA was sequenced 48, 72, and 96 h post-exposure. A DuplexSeq mutagenicity panel targeting twenty 2.4-kb regions distributed across the genome was used to sample diverse, genome-representative sequence contexts. A robust increase in MF that was unaffected by time was observed in both laboratories. Concentration-response in the MF from the two laboratories was strongly positively correlated (R^2^=0.95). C:G>T:A, T:A>C:G, T:A>A:T, and T:A>G:C mutations increased in consistent, concentration-dependent manners in both laboratories, with high proportions of C:G>T:A at all time points. The target sites responded similarly between the two laboratories and revealed a higher average MF in intergenic regions. These results, demonstrating remarkable reproducibility across time and laboratory for both MF and spectrum, support the high value of DuplexSeq for characterizing chemical mutagenicity in both research and regulatory evaluation.

## INTRODUCTION

Mutagenicity testing is an integral part of chemical and pharmaceutical toxicity assessment because mutations can cause cancer, age-related conditions, and genetic diseases. The current standard *in vitro* mutagenicity assays in mammalian cells (*e*.*g*., *HPRT, XPRT, or Tk+/-* gene mutation assays^1, 2^) determine the frequency of mutations in specific reporter genes. Together with the Ames test in bacteria^3^, these assays have been effective for mutagenic hazard identification and served as gold standards in genetic toxicology for decades. However, it is increasingly recognized that the current standard *in vitro* assays provide an incomplete picture of the genome-wide effects of mutagens.

The *in vitro* mammalian cell gene mutation assays are restricted to the analysis of simple mutant frequency in single, sometimes exogenous, reporter genes. They also provide limited information on the specific types of mutations needed to infer mechanisms of mutagenicity without extensive work on individual mutant clones. Importantly, these methodologies are limited to specific *in vitro* models and cannot readily be integrated with other model systems and assays. Consequently, there is a need for direct, quantitative mutation detection methods for mammalian genomes that readily provide information on mutational spectrum to improve mutagen identification and mechanistic understanding.

Next-generation sequencing (NGS) enables quantitative analysis of mutation spectra and its application in genetic toxicology is becoming more prevalent. NGS can provide information beyond simple mutant frequency by quantifying each type of base substitution and more complex forms of mutation, as well as identifying the local sequence context in which the mutations occur across the organism’s genome^4, 5^. However, the high per base pair rate of technical errors (on the order of 1 in 10^3^) of NGS leads to an overestimation of the true MF in cells that has hindered its application.

To circumvent this problem, error-corrected sequencing (ecNGS) techniques have been developed to improve accuracy and specificity for detecting true, low frequency mutations^6-8^. The highest resolution of ecNGS methods, such as Duplex Sequencing (DuplexSeq; TwinStrand Biosciences, Seattle, WA), involves labeling both strands of a DNA duplex with one or more forms of a unique molecular index (UMI) that both relates and distinguishes the sequences of the two strands from each other and from those of other molecules. Following sequencing, this information is used to create a consensus sequence that eliminates errors introduced during DNA handling, PCR amplification, and sequencing. DuplexSeq improves the error rate of conventional NGS by approximately 10,000-fold, enabling the detection of a mutation present among 10^7^ bases or more^6, 9-11^.

The DuplexSeq approach enables the quantification of locus-specific mutation frequency (MF) and quantitative measurements of the extent of clonal expansion of mutations when applied in a targeted form using hybrid selection panels^12^. Genome-representative panels have been designed specifically for mutation analysis in human, mouse, or rat genomes for chemical mutagenicity assessment; the panels contain 20 endogenous genomic sites, each 2.4 kbp in length, randomly spread across 20 autosomal chromosomes (TwinStrand DuplexSeq Mutagenesis kits). The mutagenesis panels sample sequences that are collectively representative of the genome in terms of GC-content, genic and intergenic regions, and coding and non-coding sequences^13^. Selective sequencing at high depths makes DuplexSeq practical for higher throughput applications as a broadly accessible mutagenicity assay.

Within the field of genetic toxicology, DuplexSeq with genome-representative hybrid selection panels is an emerging methodology and only a handful of reports have focused on its application to chemical mutagenicity assessment. Valentine et al.^12^ and LeBlanc et al.^13^ demonstrated that DuplexSeq produced *in vivo* results that were comparable to, yet more data-rich than, TGR assays. Wang et al.^14^ demonstrated its ability to detect a concentration-dependent increase in mutations in a specialized *in vitro* airway tissue model comprising donor-derived primary human cells exposed for 28 days to the potent alkylating agent ethyl methanesulfonate (EMS). The results thus far demonstrate the added value of DuplexSeq in investigating chemical mutagenicity.

However, the utility, feasibility (e.g., inter-laboratory transferability and reproducibility), and optimal approach to incorporating DuplexSeq into a standardized, high-throughput *in vitro* mutagenicity assay for chemicals in regulatory testing are as-of-yet unaddressed. Advancing this application will require extensive validation studies benchmarked against traditional mutagenicity assays. Furthermore, it is necessary to investigate how the timeline and concentration selection for *in vitro* DuplexSeq mutagenicity assays can be integrated with standard genetic toxicology assays, such as the micronucleus (MN) test, to produce a modernized and efficient genotoxicity test paradigm.

The objectives of the work presented here were to: (1) identify the optimal sampling time for mutation analysis in human lymphoblastoid TK6 cells – a widely used *in vitro* model in regulatory genetic toxicology with a doubling time of approximately 12 - 14 h^15^; (2) evaluate inter-laboratory consistency of DuplexSeq results in this model; and (3) explore the additional information gained by analyzing NGS-based mutational endpoints in the 20 target sites. Toward this, we conducted two independent *in vitro* time-series and concentration-response analyses of a prototype mutagen using the same DuplexSeq methodologies at two different facilities: the Genomics Laboratory at Health Canada (HC; Ottawa, ON, CA) and Inotiv, Inc. (Inotiv-RTP; Research Triangle Park, North Carolina, US). Both laboratories independently exposed TK6 cells to *N*-ethyl-*N*-nitrosourea (ENU), a historical positive control for mutagenicity, and used viability measurements to select the top concentrations. Cells for DuplexSeq were sampled at the same time points over a 96 h period in both laboratories. DuplexSeq libraries were prepared at TwinStrand BioSciences; a replicate set of DNA for one time point was also used to independently prepare libraries at Health Canada. We examined the overall MF induced by ENU at different concentrations and time points, the absolute frequencies and relative proportions of individual base substitution types, and the concentration-response relationships to evaluate assay performance and inter-laboratory concordance in data generation. The DuplexSeq panel also enabled the comparison of MF across the target sites located in different chromosomes, as well as the trinucleotide mutation spectra. In aggregate, our findings indicate that a DuplexSeq-based approach for quantitative *in vitro* chemical mutagenesis assessment in human cells is feasible, highly reproducible across laboratories, and unlocks previously inaccessible types of genomic data relevant to genotoxicity assessment.

## MATERIALS AND METHODS

### Cell culture and exposure

Both the HC Genomics Laboratory and Inotiv-RTP independently obtained the human TK6 lymphoblastoid cells from American Type Culture Collection (ATCC) (ATCC# CRL-8015; Manassas, VA). At HC, cells were maintained in T75 flasks at a density between 1×10^5^ and 1×10^6^ cells/mL in RPMI 1640 media (Gibco) supplemented with 10% v/v heat-inactivated horse serum (Gibco; New Zealand origin), 1 mM sodium pyruvate (Gibco), and 2 mM L-glutamine (Gibco), at 37°C in 5% CO_2_. At Inotiv-RTP, TK6 cells were maintained in RPMI 1640 medium (Gibco) supplemented with 10% v/v heat-inactivated horse serum (Gibco; New Zealand origin), 1% v/v Pluronic F-68™ (Gibco), 0.5 % sodium pyruvate (Gibco), and antibiotics (penicillin at 20 Units/mL and streptomycin at 20 µg/mL) (Gibco), at 37 ± 1 °C in 6 ± 1 % CO_2_.

ENU (Sigma Aldrich) solutions were prepared in dimethyl sulfoxide (DMSO) (HC: Fisher Scientific; Inotiv-RTP: Sigma Aldrich) immediately prior to exposure at both facilities. At HC, cells were transferred to 6-well plates at a density of approximately 3 × 10^5^ cells/mL with 5 mL of cell suspension in each well. Cells were exposed to a range of ENU concentrations (25, 50, 100, and 150 µM) or an equivalent volume of DMSO (1% v/v) in three replicates (n=3) at 37 °C and 5% CO_2_. After 24 h, cells were counted and 3 mL of each cell suspension were removed, while the remaining 2 mL of the suspension were transferred to a new 6-well plate. Three mL of RPMI 1640 media (supplement composition as described above) were added to each well to a final volume of 5 mL. Cell counting and sub-culturing procedures were repeated at the 48 h and 72 h time points following the initial exposure. At the 48 h and 72 h time points, 3 mL of cell suspension were collected and pelleted, followed by washing twice in 1x PBS (pH 7.4) before freezing the cell pellets at -80°C. At the 96 h time point, all cells were collected and frozen after counting and the two PBS washes. For the MN assay, a 96-well plate was prepared with 99 µL of TK6 cell suspension in each well at a density of 3 × 10^5^ cells/mL. Cells were exposed to the same concentrations of ENU as described above with four replicates of each concentration (1% v/v). After 24 h, the MN assay was performed as described below.

At Inotiv-RTP, 50 ± 0.1 mL of TK6 cell suspension at a density of 4.0 ± 0.5 × 10^5^ cells/mL were prepared in 600 mL suspension culture flasks. Cells were exposed to five concentrations of ENU (25, 50, 100, 150, and 200 µM) or an equivalent volume of DMSO (1 % v/v) in duplicate flasks (n=2) at 37 °C and 6% CO_2_. After 24 h, 5 × 10^5^ cells were removed from the vehicle control flasks and equivalent volumes were collected from each ENU-treated flask to perform the MN assay. The remaining cells were pelleted and washed twice by resuspending in 1x PBS (pH 7.4) to remove the PBS. The cells were resuspended to a density of 3.5 ± 0.5 × 10^5^ cells/mL in RPMI 1640 media and 50 mL were returned to the culture flasks in the incubator. The remaining cells were rinsed twice with 1x PBS containing 1 mM EDTA and the pellets were flash frozen in liquid nitrogen before storing at -80°C. The above procedures were repeated at the 48 h and 72 h time points. After 96 h, all cells were collected and frozen after counting and washing twice with 1x PBS containing 1 mM EDTA.

### In vitro micronucleus flow cytometry assay

The MN frequency was measured using the *In Vitro*

MicroFlow kit (Litron Laboratories, Rochester, NY) on a MACSQuant Analyzer 10 flow cytometer (Miltenyi Biotec) at HC and a Becton-Dickinson FACSCalibur flow cytometer at Inotiv-RTP. The samples were prepared for the flow cytometric analysis according to Litron’s protocol for 96-well plates (HC) and for larger format vessels (Inotiv-RTP). At HC, for each ENU treatment concentration and vehicle control (n=4), 5000 cells were analyzed to determine the percentage of apoptotic cells, MN frequency, and the ratio of nuclei and the counting beads added to all samples during assay preparation as per the protocol. At Inotiv-RTP, 20,000 cells were analyzed from each culture replicate (n=2 for each ENU-exposure and the vehicle control) for the same parameters. Relative survival (RS) was determined by dividing the nuclei-to-bead ratio of all treated samples by that of the corresponding vehicle control. Apoptotic cells and MN were detected by the double staining method described in Litron’s protocol.

### DNA isolation and quality assessment

DNA extraction from the HC TK6 cell pellets was performed in-house in Ottawa, Canada and frozen DNA samples (two replicate sets of samples collected at 48, 72, and 96 h) were shipped to TwinStrand Biosciences on dry ice. Two replicate sets of 48 h DNA samples were retained for in-house DuplexSeq library preparation at HC. The frozen cell pellets produced by Inotiv-RTP were shipped on dry ice to TwinStrand Biosciences for DNA extraction and DuplexSeq library preparation. In both laboratories, DNA samples were extracted and purified using the DNeasy Mini Kit (Qiagen) following the manufacturer’s protocol. DNA was eluted in TElow (10 mM Tris-HCl pH 8.0, 0.1 mM EDTA). The quantity of DNA in each sample was measured using a Qubit 4 fluorometer and a dsDNA High Sensitivity Assay kit (Invitrogen). The extent of fragmentation of DNA was assessed using an Agilent TapeStation system with Genomic DNA ScreenTape reagents (Agilent Technologies). All DNA samples had a DNA integrity number (DIN) above 8.0.

### Duplex Sequencing library preparation and sequencing

DuplexSeq was performed using the TwinStrand v1.0 Human Mutagenesis panel consisting of 20 target sites that are distributed across 20 autosomal chromosomes (Supplementary Table S1). The target regions are each 2,400 base pairs in length and include genic and intergenic sequences that are representative of the complete genome with regard to % GC-content and relative proportions of coding and non-coding regions (TwinStrand Biosciences). The 48 kb target territory excludes highly repetitive elements and pseudogenes where mapping quality could be compromised. The panel excludes genes with any significant reported role in cancer based on the COSMIC database to reduce the possibility of positive or negative selection influencing induced mutations in cells^12^. Library preparation was performed at the TwinStrand facility and at HC as previously described^13, 14^. Inotiv-RTP samples and HC samples were processed and sequenced in August 2020 and in March 2021, respectively, at the TwinStrand facility. Samples were blinded during the analysis at TwinStrand. At HC, libraries for two additional replicates of 48 h samples were prepared in September 2021.

Briefly, for each sample 1000 ng of DNA was ultrasonically sheared to a median size of ~300 bp, end-repaired, A-tailed, and ligated to DuplexSeq Adapters and underwent target enrichment using the reagents and standard protocol included with the TwinStrand DuplexSeq Human Mutagenesis™ kit. The libraries prepared by TwinStrand were sequenced using 150 bp paired-end reads on an Illumina NovaSeq 6000. The 48 h libraries prepared at HC were shipped to and sequenced at Psomagen (Rockville, MD, US) under identical conditions.

TwinStrand Biosciences processed all raw sequencing data through the DuplexSeq human mutagenesis pipeline as previously described^12^. The raw sequencing data from the 48 h samples that were processed at HC were analyzed separately from the sequencing data produced by TwinStrand Biosciences. Mutation frequency (MF) is determined by dividing the number of unique mutations (i.e., ignoring the effect of clonal expansions) by the total number of duplex bases sequenced. The TwinStrand Biosciences pipeline provides sequencing quality metrics, MF, mutation spectra, trinucleotide frequency and MF per target data.

### Statistical Analysis

Statistical analysis, including the estimation of MF and pairwise comparisons to controls, were conducted in the R environment for Statistical Computing^16^. Generalized linear models were fit to the data using the glm() function assuming a binomial error distribution. The Analysis of Deviance Table was estimated using the Anova() function in the cars R package^17^. For generalized linear models, this function presents the likelihood-ratio chi-square statistic from the model as an ANOVA table. Pairwise comparisons were conducted using the doBy R package^18^. The p-values from the hypothesis tests comparing the MF at each concentration to controls were adjusted for multiple testing using the Holm-Sidak correction. This multiple testing correction was applied within each time point and laboratory. This analysis was conducted on the pooled target sites as well as at each target site independently. Generalized linear mixed models were used to analyse the differences between target sites and laboratories and adjusted for multiple testing using the Holm-Sidak correction. The Wald Test was used to compare intergenic and genic target sites.

### Trinucleotide mutation spectra and Catalogue of Somatic Mutations in Cancer (COSMIC) signature analysis

Cosine similarity between the mutational spectra obtained in the two laboratories and COSMIC signatures was evaluated in the R environment for Statistical Computing^16^ using the MutationalPatterns 3.4.0 package^19^. First, mutation counts in the two replicates of each concentration at each time point were pooled. A control signature was generated at each time point using the vehicle control spectra. Using the fit_to_signatures_strict() function, the optimal combination of mutational signatures (among the control signature and the COSMIC signatures) that had the highest cosine similarity to each ENU-treated mutation spectrum was identified and the number of mutations that contributed to each signature in the mutation spectrum was determined. In addition, the cos_sim_matrix () function was used to evaluate the cosine similarity between each COSMIC signature and each ENU-treated mutation spectrum. Using the same function, the cosine similarity between the trinucleotide spectra from the two laboratories were analyzed.

### Benchmark concentration modeling

Using the PROAST web application (version 70.1; RIVM National Institute for Public Health and the Environment; https://proastweb.rivm.nl), the concentration-response of MF was modeled using the Hill model to determine the benchmark concentration (BMC) of ENU that induced a 50% increase in MF at 48 h, 72 h, and 96 h. The limit for the AIC (Akaike Information Criterion), which describes the goodness of fit of the model, was set at 2. The lower and upper 90% confidence limits (BMCL and BMCU, respectively) were estimated in addition to the BMC.

## RESULTS

### Comparison of cytotoxicity and micronucleus frequency

The flow cytometry MN assay is a widely used assay that would fit well within an integrated test with DuplexSeq to identify potentially clastogenic and mutagenic agents. To select the concentrations for mutation analysis that induce DNA damage without excessive cell death, the MicroFlow assay was used to measure the MN frequency, relative survival (RS), and apoptotic and necrotic population in TK6 cells 24 h after exposure to a range of ENU concentrations at HC and Inotiv-RTP (Figure 1). These measurements also provided an additional point of comparison to assess the consistency in exposure in the two laboratories.

**Figure 1.**
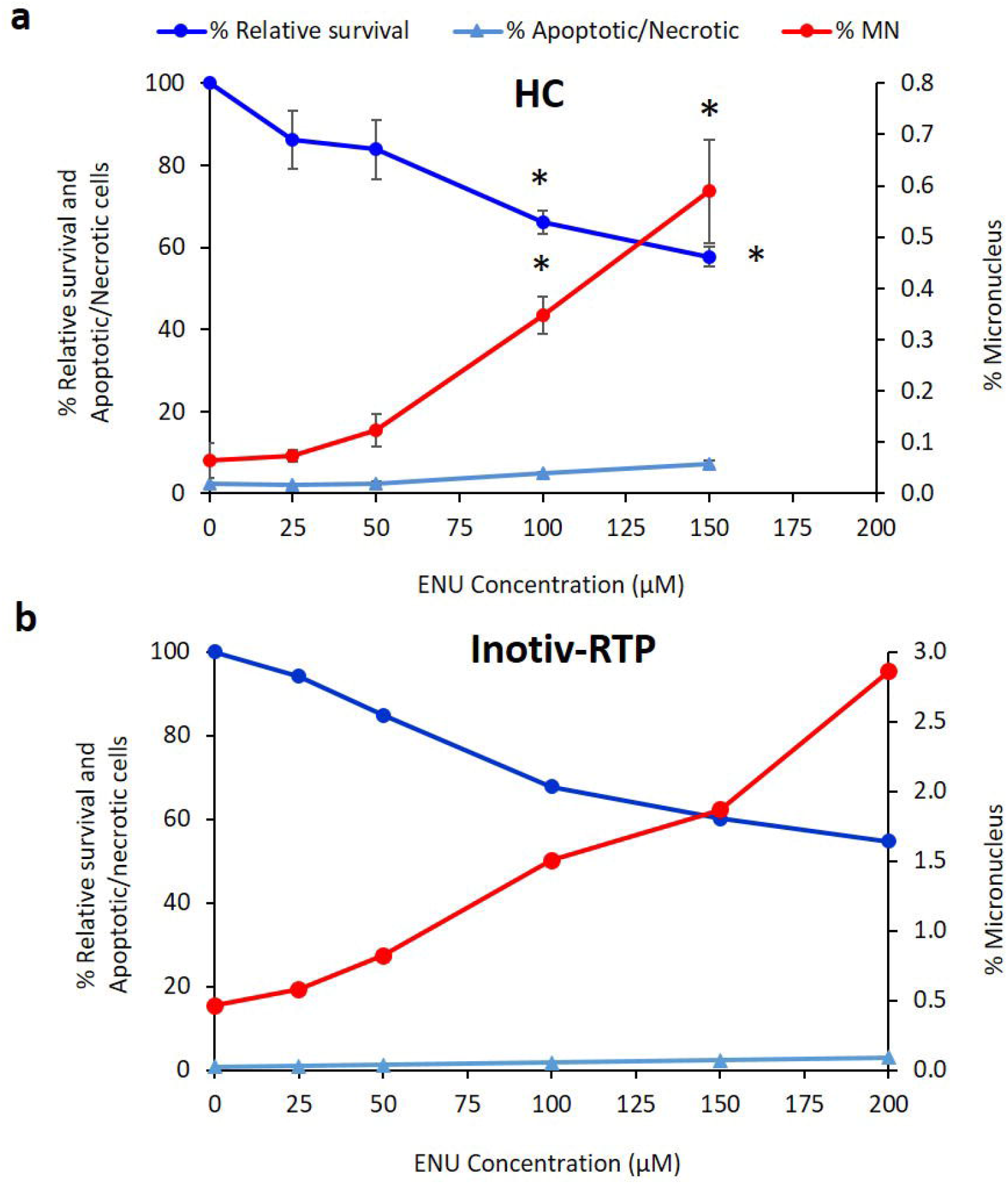
Micronucleus measurement in TK6 cells exposed to ENU in two different laboratories, HC (top) and Inotiv-RTP (bottom), using the In Vitro MicroFlow® assay (Litron Laboratories) on a flow cytometer. Percentage micronucleus was measured 24h after exposure (HC: n=4; Inotiv-RTP: n=2). Relative survival was measured as the ratio between nuclei and counting beads relative to that of the vehicle control. The error bars in the HC plot represent standard error. Asterisk (*) indicates p-value<0.0001 in one-way ANOVA with post-hoc Dunnett’s test.

ENU caused a similar reduction in cell survival in the two independently performed experiments. For example, at 150 µM, the highest concentration used at HC and the second highest at Inotiv-RTP, the average cell survival relative to the vehicle control was 57% and 60%, respectively. At HC, the 150 µM exposure induced a 3.1-fold increase in the apoptotic and necrotic cell population, while at Inotiv-RTP, 150 µM induced a 2.2-fold increase.

In both laboratories, ENU induced concentration-dependent increases in % MN. Table 1 summarizes the average fold change in MN frequency at each ENU treatment concentration compared to the vehicle control. The average % MN in the vehicle control in the HC experiment was 0.064%. The 100 µM exposure caused a 5.4-fold increase compared to control, increasing to 0.35% MN. In the Inotiv-RTP experiment, the average % MN in the vehicle control was 0.46%, which increased by 3.2-fold to 1.5% MN at 100 µM. The 100 µM concentrations led to an approximate 32-34% decline in relative survival in both laboratories. Concentrations larger than 100 µM were excluded from the HC study because the relative increase in cell count (RICC) at 100 µM was approximately 50% after 24 h and dropped below 0% at 48 h at HC; thus, 100 µM was selected as the top concentration for analysis and comparison of the two laboratories. The concentration-response in % MN in the two experiments was strongly and positively correlated (R^2^ = 0.96) (Supplementary Figure 1). Collectively, the MicroFlow assay confirmed the comparable responses to ENU in TK6 cells in both laboratories.

**Table 1.**
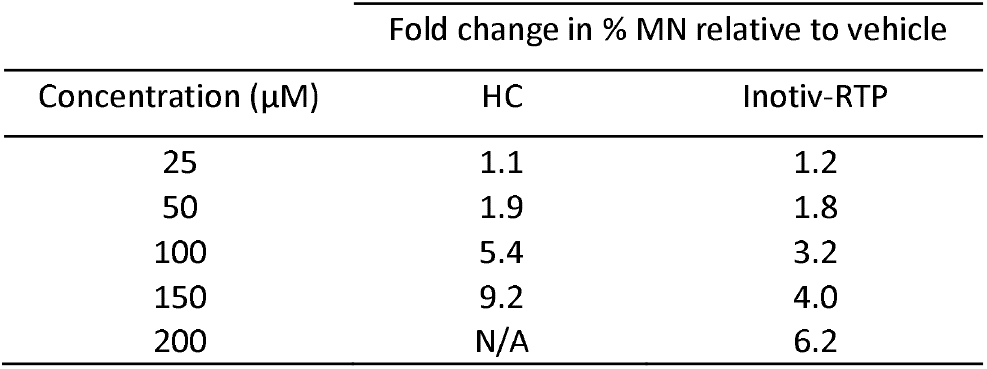
Average fold change in % MN relative to control at each ENU concentration

| Concentration (μM) | Fold change in % MN relative to vehicle | |
| --- | --- | --- |
|  | HC | Inotiv-RTP |
| 25 | 1.1 | 1.2 |
| 50 | 1.9 | 1.8 |
| 100 | 5.4 | 3.2 |
| 150 | 9.2 | 4.0 |
| 200 | N/A | 6.2 |

### DuplexSeq mutation analysis

#### 1.1. Analysis of overall mutation frequency

Unique mutations were identified by DuplexSeq in each sample and MF were calculated. Figure 2A and Supplementary Table S2 show the average MF of the two biological replicates for 0, 25, 50, and 100 µM ENU at 48 h, 72 h, and 96 h from both studies. Overall, there was a concentration-dependent increase in MF at all three time points in both datasets (Figure 2A and Supplementary Figures S2 and S3) but no significant time-dependent trend in MF at any of the concentrations in either laboratory.

**Figure 2.**
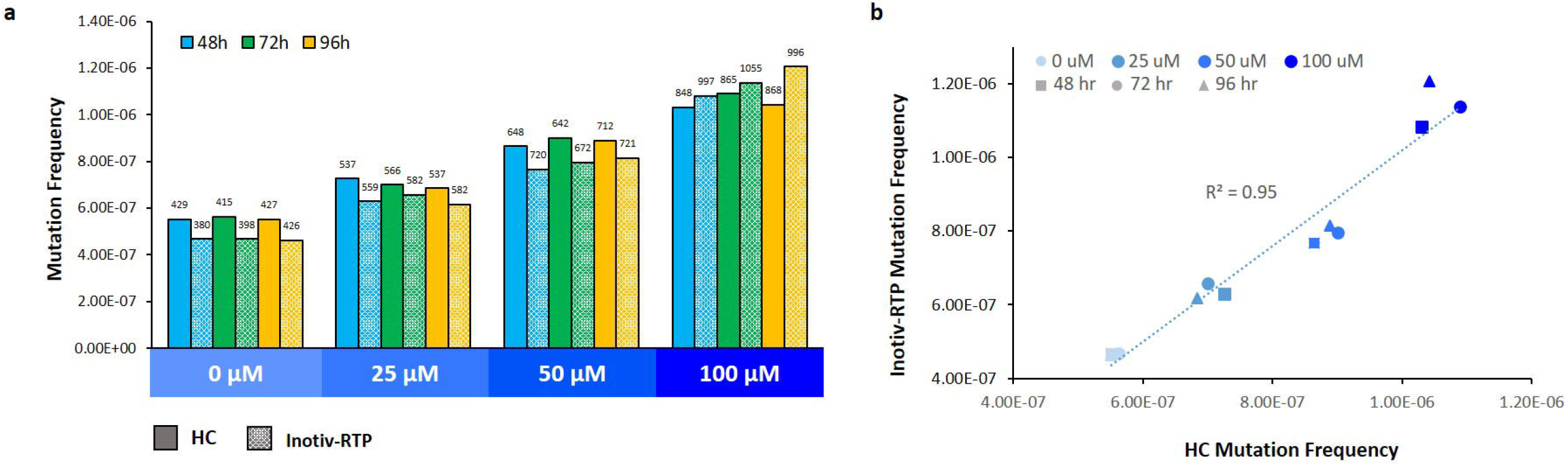
A. Mutation frequency in TK6 cells exposed to ENU measured by Duplex Sequencing (TwinStrand Biosciences). B. Linear correlation of mutation frequency measured at HC and Inotiv-RTP. Cells were sampled after 48, 72, and 96 h following the initial exposure (n=2/ concentration and time point). Library construction and sequencing were completed by TwinStrand. Numbers above the bars represent non-normalized mutation counts. Frequency was determined by dividing the counts by the total number of BP sequenced. There was a significant concentration main effect (Holm-Sidak adjusted p-value < 0.0001) and concentration by lab interaction (Holm-Sidak adjusted p-value < 0.0001). There was no statistically significant difference in mutation frequency across the three time points

The overall MF from the HC and Inotiv-RTP studies were strongly positively correlated (R^2^ = 0.95) (Figure 2B). At HC, the median MF in the vehicle control was approximately 5.5 × 10^−7^ and increased by an average of 1.9-fold at 100 µM to a median of 10.4 × 10^−7^. At Inotiv-RTP, the MF increased from a median of 4.7 × 10^−7^ in the vehicle control to a median of 11.4 × 10^−7^ at 100 µM, resulting in a 2.5-fold change. In the replicate set of 48 h samples where the libraries were prepared at HC, the average MF was 6.8 × 10^−7^ in the vehicle control and 12.0 × 10^−7^ at 100 µM, resulting in a 1.8-fold increase in MF (Supplementary Figure S2). The MF and the fold change in MF were congruent across the three datasets.

To further confirm that concentration-response to ENU in the overall MF was consistent in both laboratories at all three time points, benchmark concentration (BMC) modeling was applied. The BMC_50_ values were determined using the Hill model (Figure 3 and Table 2); a benchmark response (BMR) of 50% is recommended for various mutagenicity assays and was, thus, selected as a point of comparison across the three time points ^20^. All HC BMC_50_ 90% CIs overlapped with the Inotiv-RTP values, indicating that there were no significant differences in concentration-response across the time points and between the two laboratories.

**Figure 3.**
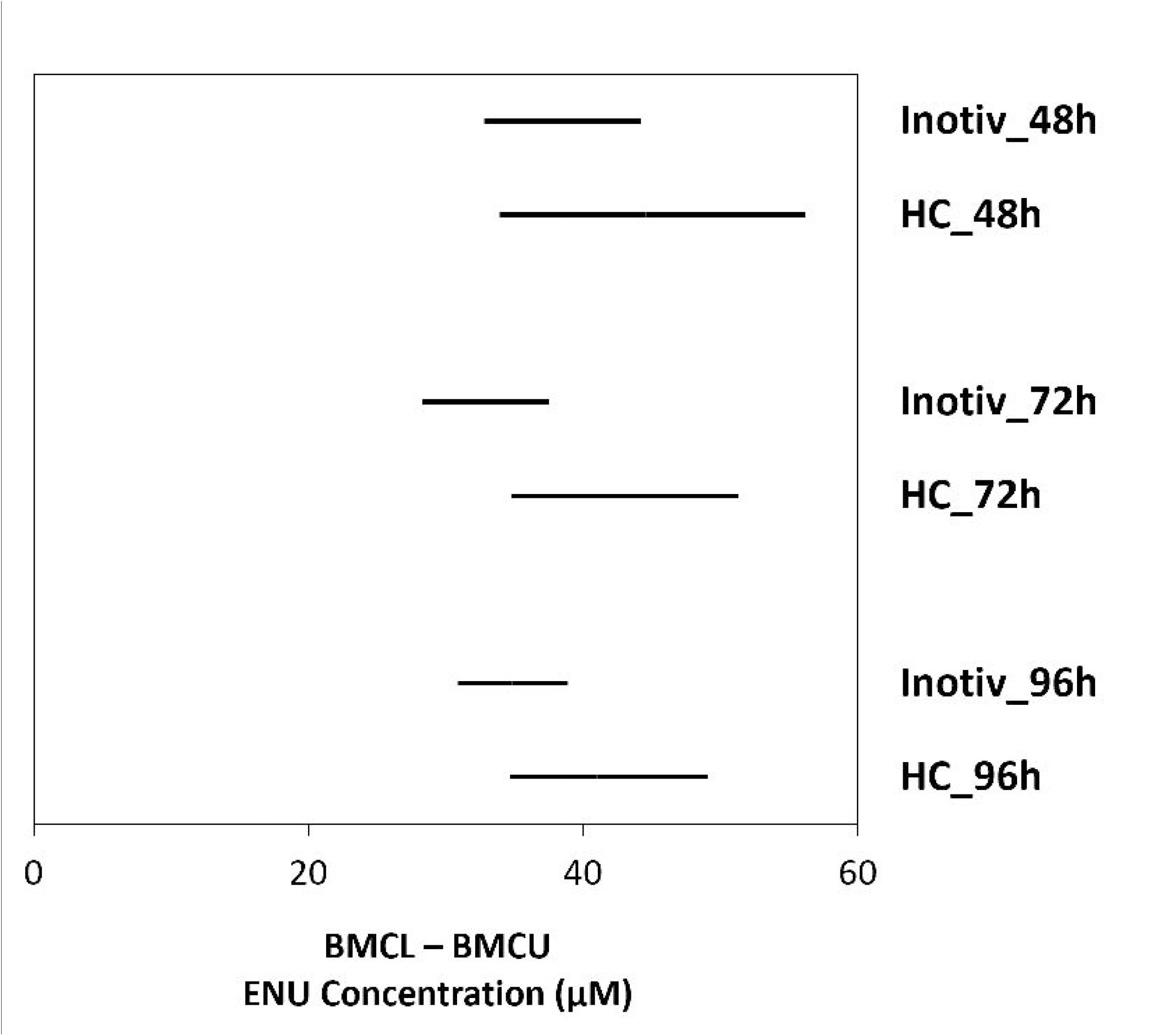
Benchmark concentrations (BMC50) of ENU that induced a 50% increase in mutation frequency 48, 72, and 96h following the initial exposure in TK6 cells at HC and Inotiv-RTP. BMC modeling was performed using the Hill model. The bars represent the 90% confidence interval of BMC50.

**Table 2.** Benchmark concentration (BMC; µM) that induced 50% increase in mutation frequency above control.

| Hill model $\text{BMC}_{50}$ (BMCL, BMCU) | | |
| --- | --- | --- |
| Individual analysis |  |  |
|  | Inotiv-RTP | HC |
| 48h | 38.34 (32.8, 44.2) | 44.55 (33.9, 56.2) |
| 72h | 33.26 (28.3, 37.5) | 41.98 (34.8, 51.3) |
| 96h | 34.84 (30.9, 38.9) | 41 (34.7, 49.1) |

### Mutation frequency across target regions

Typically, *in vitro* mutation assays are limited to detecting mutations in a single reporter gene. In contrast, the DuplexSeq Human Mutagenesis panel samples sequences from 20 different autosomal chromosomes. The panel includes 10 target regions that contain only genic sequences, six that are intergenic, and four that overlap both intergenic and genic sequences. We examined the MF in the individual target sites to determine if there were variabilities in the effects of ENU exposure in different regions of the genome. In addition, we examined the concordance in response at the 20 targets sites between the two laboratories.

We first examined the background MF in each target region by averaging the MF in the vehicle control across 48 h, 72 h, and 96 h because there was no statistically significant difference in the three time points; we then rank ordered the target regions by MF (Figure 4 and Supplementary Figure S4). Four of the five regions with the lowest background MF were the same in both laboratories (chromosomes 1, 6, 13, and 14). The two laboratories also shared four targets in the top five of the ranking (chromosomes 2, 12, 16, and 19). In the HC study, the difference in MF between the target site with the highest baseline MF (chromosome 21) and the lowest (chromosome 14) was 3-fold. In the Inotiv-RTP study, the difference in MF between the target site of the highest baseline MF (chromosome 9) and of the lowest (chromosome 14) was 2.6-fold.

**Figure 4.**
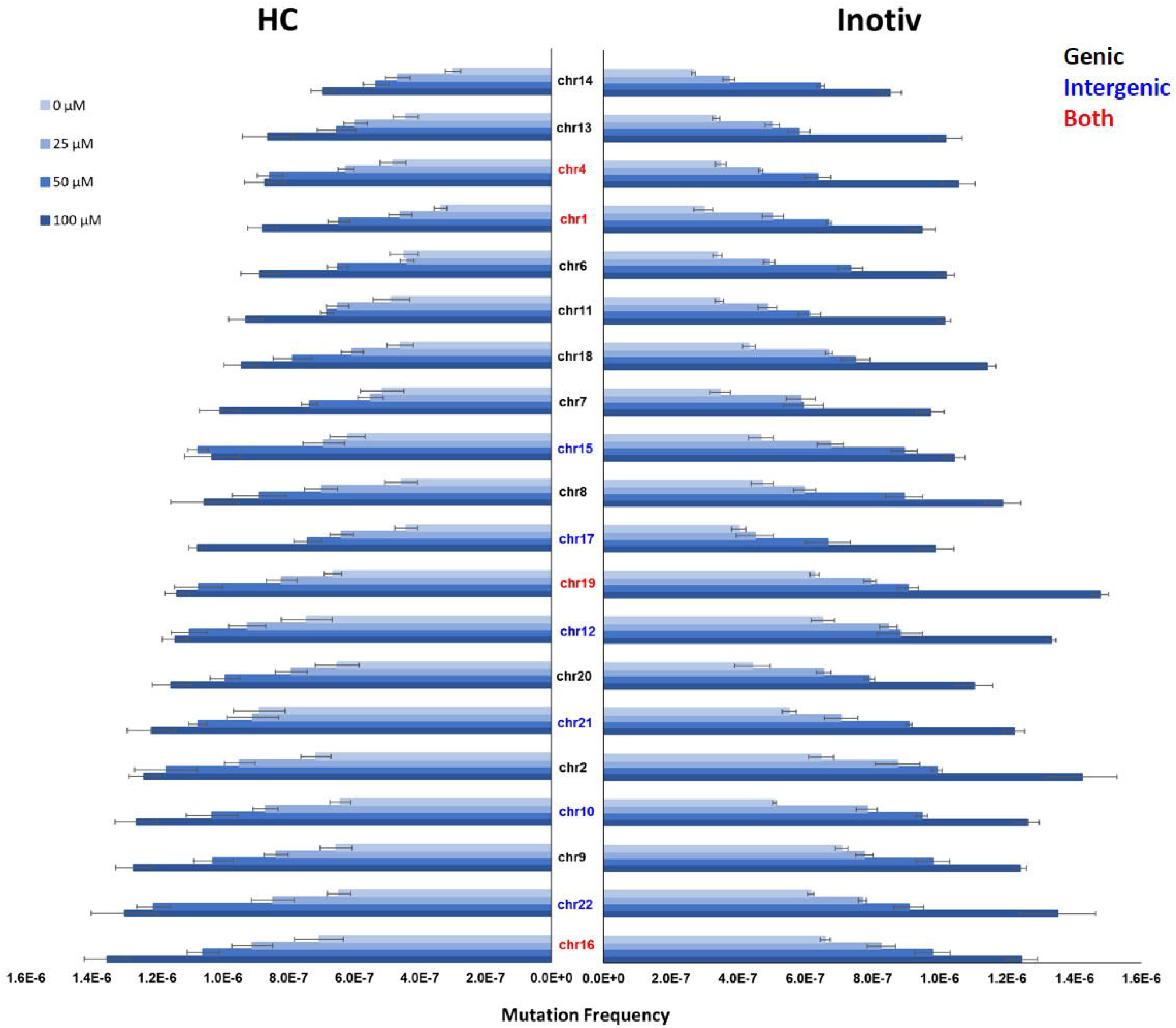
Mutation frequency in the 20 target sites of the Duplex Sequencing human mutagenicity panel measured in TK6 cells exposed to ENU at HC (left) and Inotiv-RTP (right). The mutation frequency at 48, 72, and 96 h were averaged (n=2 at each time point; n=6 in total). The target sites are listed on the Y-axis. Black indicates genic sites, blue indicates intergenic sites, and red indicates those that include both genic and intergenic sequences. The target sites are ordered according to mutation frequency in HC samples at 100 µM from the lowest (top) to highest (bottom). Error bars represent standard error.

Next, we explored target responses to ENU. There was an observable concentration-dependent increase in MF at all 20 target sites in both studies (Figure 4). After ranking the targets by MF at 100 µM, we compared the target sites in the top and bottom 5 of the ranking for HC and Inotiv-RTP. The target sites on chromosomes 2, 10, and 22 were present in the top 5 for both laboratories. In the bottom 5, the chromosomes 14 and 1 target sites were shared by the two laboratories. In the HC study, the difference in MF between the highest ranked (chromosome 16) and the lowest ranked (chromosome 14) target sites was 1.9-fold. In the Inotiv-RTP study, the chromosome 19 target ranked the highest and the chromosome 14 target ranked lowest, with a difference of 1.7-fold between the two. On average, there was a 30% increase in MF in the intergenic target sites compared to the genic sites.

### Analysis of interactions between experimental parameters

In order to determine if there were any statistically significant effects of laboratory (HC vs. Inotiv-RTP) on MF across the target sites at each concentration, we applied a generalized linear mixed model (GLMM) followed by the Holm-Sidak multiple testing correction to identify interactions between these two variables. There were no significant interactions between laboratory and target site at any of the concentrations, indicating that all loci responded similarly between the two laboratories. The interaction plots of MF across the 20 target sites at each ENU concentration in the two laboratories showed approximately parallel lines for the two datasets Supplementary Figure S5a-d).

Next, the effects of concentration, laboratory, and time point on the overall MF (pooled analysis) as well as on MF at each target site and the interaction between these experimental parameters were examined in an Analysis of Deviance Table (Supplementary Tables S3 and S4). The Likelihood Ratio (LR) test statistic indicated highly significant and prominent concentration-related effects on the overall MF and at all 20 target sites. There was no evidence of a time point-related effects on the overall MF and only one target site was significantly affected by time point. In terms of the influence of laboratory on MF, the pooled analysis indicated significant effects. However, upon analyzing individual target sites, only three target sites showed significant laboratory-related effects on MF, while at only two sites, an interaction between laboratory and concentration occurred at the highest exposure concentration. All in all, there were strong consistencies between the two laboratories across the 20 target sites.

### Mutation spectra analysis

To explore the mechanistic information produced by DuplexSeq, and to compare the consistency of the mutation spectra across the two laboratories, MF were analyzed by base substitution type (Figure 5). As described above, the replicates across the three time points were averaged for this analysis. In both the HC and Inotiv-RTP control samples, C:G > T:A was the predominant type of mutation in the vehicle control followed by C:G > A:T with similar frequencies and proportions in the two laboratories. Overall, both the proportions and frequencies of C:G > T:A, T:A > C:G, and T:A > A:T significantly increased with ENU concentration by highly similar magnitudes in the two laboratories.

**Figure 5.**
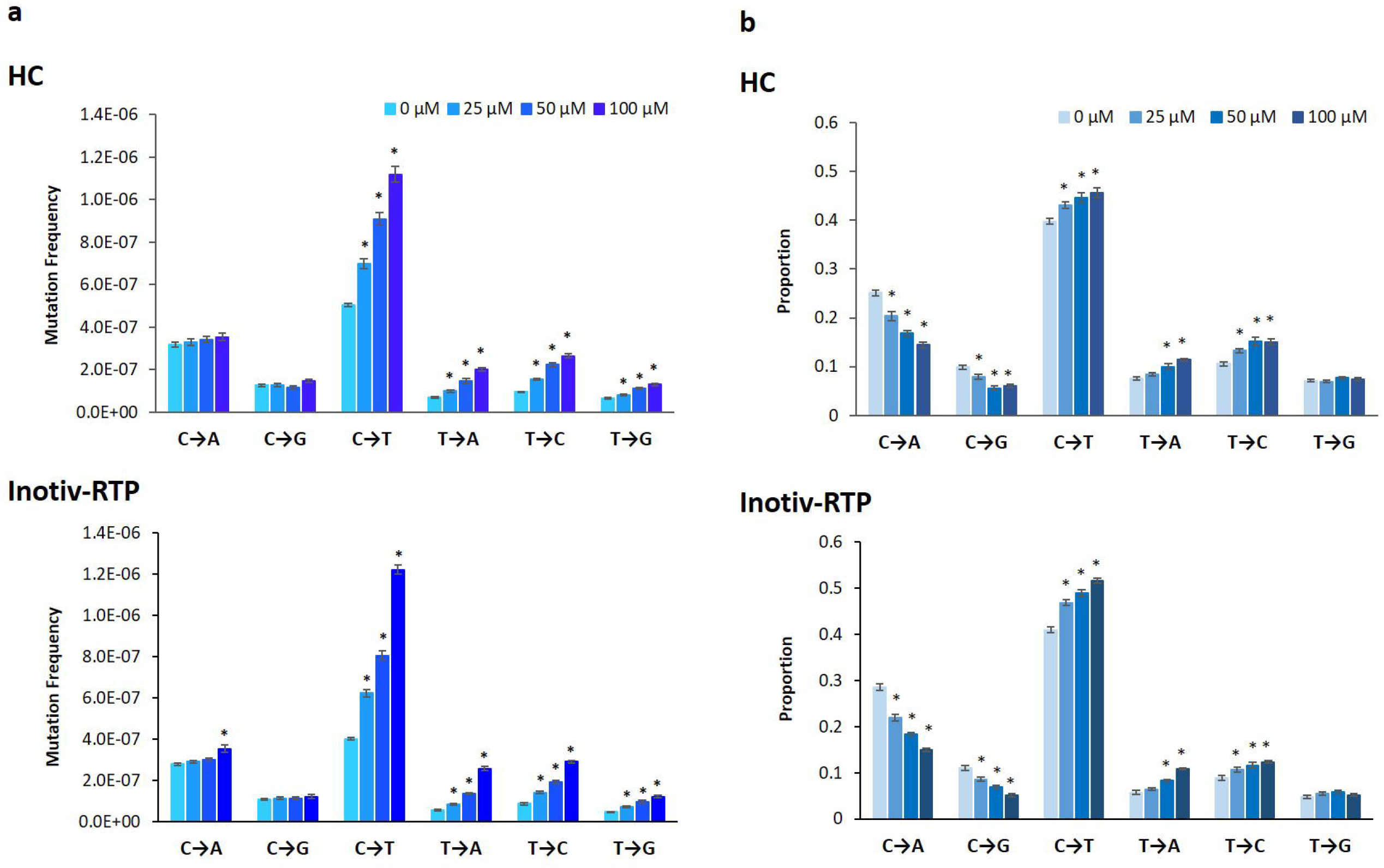
Frequencies (A) and proportions (B) of individual base substitution types in TK6 cells exposed to ENU at HC and Inotiv-RTP measured by Duplex Sequencing (TwinStrand Biosciences). Cells were sampled 48, 72, and 96 h following the initial exposure (n=2). The mutation proportions and frequency at the three time points were averaged (n=6 in total). The error bars represent standard error and asterisk (*) indicates statistical significance (p-value <0.05) in ANOVA with post-hoc Dunnett’s test.

The mutation spectra data were examined in more detail by analyzing base substitutions in each trinucleotide context as a trinucleotide spectrum (Figure 6). Consistent with all prior analyses, there was no time-dependent change in the spectra. Concentration-dependent changes in the trinucleotide spectra were observed in the COSMIC signature analysis (Supplementary Figure S6), with increases in the proportions of SBS7a and b (UV exposure-related) and SBS32 (azathioprine exposure /immunosuppression) contributing to the overall ENU mutational spectra in TK6 cells. In addition, the cosine similarity of the ENU spectra to SBS7a and b, SBS11 (temozolomide exposure), SBS30 (defective BER due to NTHL1 deficiency), and SBS32 also increased with concentration at all three time points in both laboratories (Supplementary Table S5). Overall, the two laboratories produced concordant results in the COSMIC analysis of trinucleotide spectra, indicating a high level of consistency in the mutation spectrum data produced by DuplexSeq.

**Figure 6.**
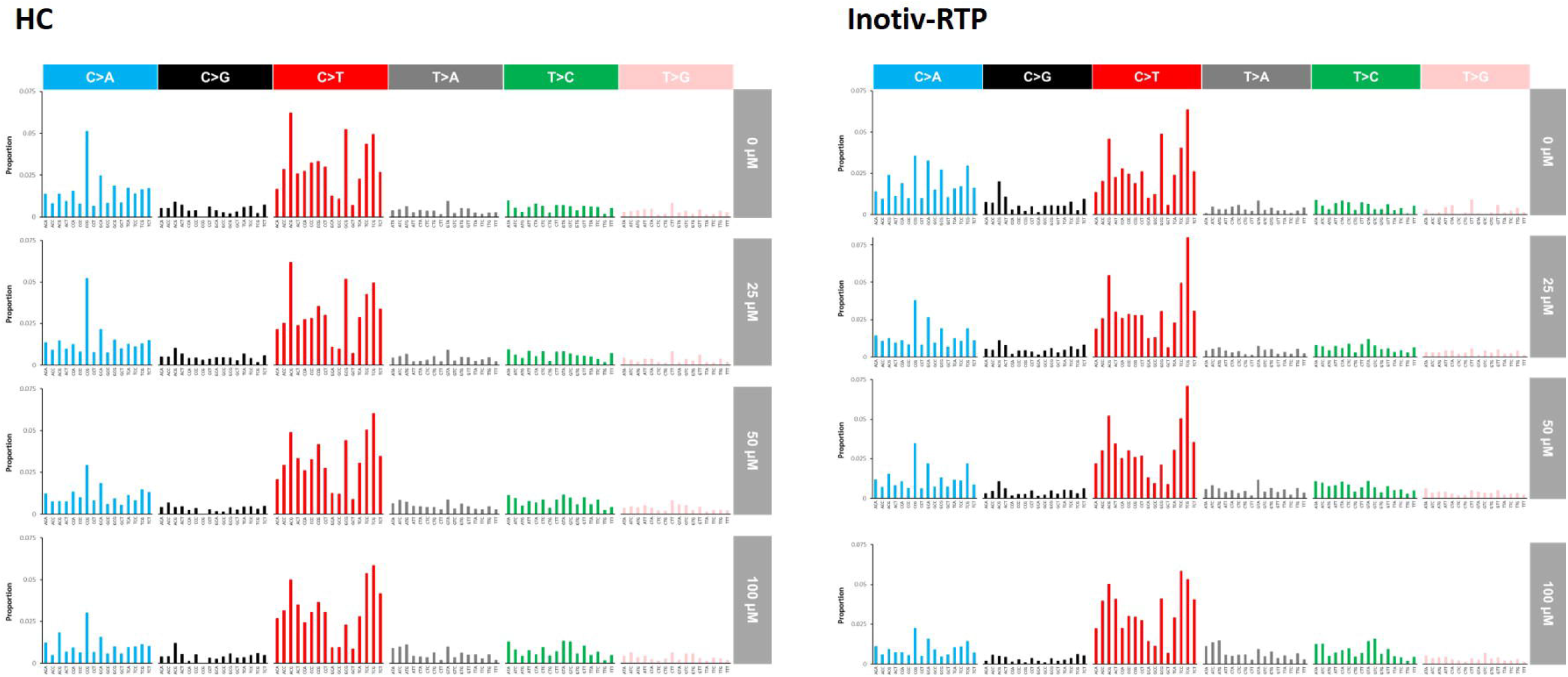
Trinucleotide mutation spectra in TK6 cells exposed to ENU at HC and Inotiv-RTP. Mutation frequency measured at 48, 72, and 96 h were averaged for each concentration.

## DISCUSSION

Our study demonstrates how error-corrected DuplexSeq of a 48-kb panel of genic and intergenic regions within the human genome can precisely and reproducibly quantify *in vitro* MF for genetic toxicology applications. The human mutagenicity panel, which contains 20 targets across different autosomes, was specifically developed for applications in chemical and pharmaceutical mutagenicity assessment. This panel-based approach provides a pragmatic way to generate sufficient sequencing data for determining locus-specific MF and to perform an in-depth investigation into the characteristics of mutagenicity by analyzing the mutation spectra. The methodology is remarkably robust, demonstrating highly concordant results across time points and between sample preparations in different laboratories. Moreover, the results acquired provide detailed mechanistic insight enabling the understanding of target nucleotides and region-specific effects of mutagens.

The first objective of this comparative study was to determine an appropriate time point for analyzing mutations by DuplexSeq when assessing potentially mutagenic exposures in TK6 cells. Two laboratories, HC and Inotiv-RTP, independently exposed TK6 cells to ENU and sampled the cells 48, 72, and 96 h post-exposure to allow fixation of DNA damage into mutations. In both laboratories, the overall MF and the induced frequencies and proportions of base substitution types were highly similar for cells sampled from 48 to 96 h following the initial exposure to ENU at all concentrations. Furthermore, the overlapping BMC_50_ CIs at all sampling times in both laboratories supported the lack of time-dependent effects on concentration-response. Since the increases in MF in ENU-treated samples were statistically significant at 48 h and did not increase with longer incubations, the two later sampling times may not be necessary to detect mutations by DuplexSeq. As a point of contrast, the human TK6 cell HPRT gene mutation assay, one of the standard in vitro mutagenicity tests, takes up to 4 weeks before mutant scoring^21^. With a 48 h sampling time, induced mutation analysis by DuplexSeq drastically reduces the time and labour required compared to conventional tests, providing both an efficient and sensitive mutagenicity assessment methodology; this provides a tremendous advantage for the use of DuplexSeq for mutagenicity testing.

At Inotiv-RTP, a 24 h sampling time was also included in the experiment alongside MN analysis, with additional ENU concentrations of 150 and 200 µM at both 24 h and 96 h. While a concentration-response in the overall MF was observable at 24 h, the magnitude of response was smaller than at 48, 72, and 96 h, and the discrepancy became larger with increasing ENU concentration. This difference in concentration-response was also evident in the BMC_50_ at 24 h (52.2 µM; BMCL: 44.0 µM; BMCU: 61.3 µM), which was higher than the BMC_50_ at the three later time points measured at Inotiv-RTP. It is possible that cytostatic effects accompanying the increase in ENU concentration are causing a proportion of cells that retain ethyl adducts to remain viable but undivided at 24 h. Indeed, we observed a concentration-dependent reduction in the relative increase in cell count at 24 h, but the percentage of viable cells measured by trypan blue exclusion was unaffected up to 200 µM in our preliminary range finding experiment at HC at 24 h (data not shown). Based on these results, measuring both MN and mutations 24 h following the initial exposure is not recommended and, thus, these assays will require distinct time points for genotoxicity assessment.

Objective two aimed to assess the inter-laboratory reproducibility of mutagenicity assessment by DuplexSeq. The concordance in concentration-response in two experiments run at distinct laboratories was evaluated by correlation analysis of MF at each concentration and time point, in addition to BMC modeling at all three time points. We found a high, positive correlation between the two laboratories, with an R^2^ value of 0.95 in MF. This concordance in concentration-response was supported by the BMC analysis at each time point, where the BMC_50_ CIs derived from the two studies overlapped with each other. These finding suggest that different laboratories will obtain similar mutagenicity points of departure in DuplexSeq applications that use TK6 cells, which is critical to international efforts to adopt mutagenicity as an endpoint of regulatory concern^20, 22^

As an added support to the inter-laboratory validation, two replicate cell samples that were collected 48 h post-exposure in the same experiment at HC were retained for in-house DuplexSeq library preparation and sequenced at a third party NGS facility (Psomagen Inc., Maryland). These samples were prepared and examined independently of the HC and Inotiv-RTP samples that were shipped to and processed by TwinStrand. While the MF measured in the 48 h samples processed in-house was higher than in both the 48 h HC and Inotiv-RTP samples that were processed by TwinStrand at all concentrations, the magnitude of increase in MF was highly similar. For example, at 100 µM, MF increased by 1.8-fold compared to control in the HC-processed samples and by 1.9- and 2.5-fold in the TwinStrand-processed samples. These additional data not only demonstrate the inter-laboratory consistency of DuplexSeq between HC and Inotiv-RTP, but also the promising reproducibility of DuplexSeq even when libraries are constructed and sequenced at different facilities, months apart.

In addition to inter-laboratory concordance in MF, we evaluated whether the mutation spectra, both simple and trinucleotide, were consistent between the two laboratories. Both types of spectra remained constant over time at both HC and Inotiv-RTP and were consistent between the two laboratories across concentrations. The COSMIC signature analysis of the trinucleotide spectra revealed concentration-dependent increases in cosine similarity to the same COSMIC signatures in both laboratories at all time points. A more in-depth discussion on the observed COSMIC signatures is in the Supplementary Discussion. Overall, the analysis supports that DuplexSeq generates highly reproducible spectral data.

As our third objective, we examined the MF at each of the 20 target sites of the mutagenesis panel and investigated whether the MF at each locus was affected differently by experimental parameters including ENU concentration, laboratory, and time point. Almost all loci showed significant concentration responses in MF in both laboratories. At five target sites, there appeared to be laboratory-driven effects on MF in the analysis of deviance. The main effects at two sites were inconclusive, while the small but significant laboratory-driven effects on chromosomes 17, 20, and 21 indicated differences between the two laboratories at these three sites. This is likely due the lower baseline MF in the Inotiv-RTP relative to HC samples leading to a greater response in these target sites at Inotiv-RTP. However, we caution over-interpretation of these results because of the small sample size (n=2) and the fact that MF at these sites were within one order of magnitude with each other. Ranking of the target sites by MF in the vehicle control and at 100 µM (averaged across time points) revealed some shared target sites in the top and bottom five between the two laboratories. Furthermore, in the GLMM analysis of individual concentration groups, we observed no significant interaction between target sites and laboratory at any of the concentrations and, thus, no laboratory-dependent effects on target sites in this model. All in all, the target sites responded to ENU in a consistent manner and there was a good degree of similarity in locus-specific MF in the two different laboratories.

We further investigated what additional biological and mechanistic understanding we could gain from the mutagenesis panel that add value to mutagenicity assessment. First, we examined the differences in MF between coding and non-coding regions. We expected genic regions, which are more likely to be repaired by transcription-coupled nucleotide excision repair (TCR-NER), to contain fewer background mutations and to show less mutagenesis in ENU-exposed cells ^12, 13, 23^. Indeed, on average, the MF in the intergenic regions was 30% higher than in the genic regions, indicating differences between the two types of target regions as previously observed in an *in vivo* DuplexSeq study in transgenic rodents^13^. To estimate the transcription levels of the genic regions, we obtained transcript levels for 6 of the genic target sites from in-house DNA microarray gene expression profiles of TK6 cells exposed to a range of ENU concentrations for 4 h. These genes were all expressed at levels below the median expression level of the whole transcriptome and were not responsive to ENU treatment (data not shown). Given this, it may be possible that the genes in the genic target sites are not significantly affected by TC-NER, compared to more actively transcribed genes. Nonetheless, the work demonstrates additional biological insight that can be acquired from the use of locus-specific panels in DuplexSeq assays.

Notably, we observed an ENU mutation spectrum different from the previously reported mutation spectra in human induced pluripotent stem cell-line (iPSC)^24^ and in transgenic rodent tissues (Big Blue and TgrasH2 mice)^12^. In an iPSC model, T:A > C:G was predominant, while in TGR models, T:A > A:T and T:A > C:G were the predominant types of base substitution following ENU exposures. However, our observations are similar to the ENU-induced spectrum generated in TK6 cells using another ecNGS method, whole genome paired-end and complementary consensus sequencing (PECC-seq)^8^. The difference in mutation spectra from other models is likely due to the elevated baseline levels of C:G > T:A and C:G > A:T in TK6 cells, which accounted for 40% and 25-28% of the overall baseline mutations, respectively, at HC and Inotiv-RTP. Mutational spectra are influenced, not only by exposure to mutagens, but also by endogenous factors such as tissue-specific DNA repair mechanisms and polymerases^23, 25^, physiologic processes such as autoimmunity or inflammation^26^, inherited genetic factors, and aging^27, 28^. Indeed, the spectra in TK6 cells aligned with the known characteristics of the cell line and accompanying culture conditions; the reduced repair capacity for alkylated guanine in TK6 cells due to its MGMT deficiency paired with spontaneous deamination occurring over the course of culturing are the likely causes of the elevated levels of C:G > T:A^29-31^. A possible cause of the higher baseline frequency of C:G > A:T is oxidative damage of guanine bases forming mutagenic lesions such as 8-oxo-2’ -deoxyguanine (8-oxo-dG). Culture conditions (e.g., culture media, O_2_ level, atmospheric pressure) could increase cellular susceptibility to oxidative stress by stimulating reactive oxygen species production and/or interfering with antioxidant defences^32^. Unrepaired 8-oxodG can pair with adenine during replication and lead to C:G > A:T transversions^33^. This may lead to the accumulation of C:G > T:A and C:G > A:T with successive cell divisions. Overall, we speculate that the above factors lead to TK6 cell line-specific mutation spectra.

In summary, we identified 48 h post-exposure to be an optimal sampling time for *in vitro* mutation analysis in TK6 cells for a potent alkylating agent. Compared to gene mutation assays that are based on mutant selection and quantification, DuplexSeq, especially when combined with a genome-representative hybrid selection panel, presents a significant advantage in terms of efficiency and the diversity of information that can be obtained. However, it must be noted that the current associated costs may limit its accessibility. We also demonstrated high inter-laboratory consistency of the DuplexSeq human mutagenicity assay in two different laboratories. The results for both MF and spectra were remarkably concordant between the two experiments in all comparisons, suggesting robust reproducibility of the assay and its strong potential for test guideline development. In addition, the targeted panel approach enabled analyzing and comparing MF in different genomic contexts. Additional validation and data are required to support this emerging approach to mutagenicity testing in gaining regulatory acceptance, including its capability to detect mutagens with varying potencies and more complex mechanisms of action. Nonetheless, studies such as ours provide a step forward in building confidence in DuplexSeq for applications in genetic toxicology and set the stage for applying this methodology to modernize chemical risk assessment.

## Supporting information

Supplementary Discussion

## ACKNOWLEDGMENTS

This study was funded by Health Canada’s Genomics Research and Development Initiative (GRDI) and genetic toxicity testing conducted for the Division of Translational Toxicology (DTT), National Institutes of Environmental Health Sciences, National Institutes of Health, US Department of Health and Human Services under contract HHSN273201300009C. CLY acknowledges the Canada Research Chairs Program. We would like to thank Annette Dodge (University of Ottawa), Danielle LeBlanc, Dr. Matthew Meier (Health Canada), and Mark Consugar (TwinStrand Biosciences) for their support in conducting the DuplexSeq assay and analysis. The authors would also like to thank Dr. Scott Auerbach and Ms. Julie Foley at the DTT, and Drs. Marc Beal and Julie Cox at HC for their careful review of the manuscript.

## AUTHOR CONTRIBUTIONS

EC, CDS, and MR conducted the cell culturing, chemical exposures, and DNA sample preparation. SLSR and KLW designed the study conducted at the Inotiv facility and acquired funding from the NIEHS. EKS performed the Duplex Sequencing library building. JY, THS, PV, FYL, CCV, and JJS processed the sequencing data and performed bioinformatics analyses. AW performed all statistical analyses. CLY, FM, LR, KLW, and SLSR obtained funding to support the project. EC and CLY wrote the main manuscript text. All authors provided intellectual and technical input on the study design, data analysis and interpretation, and manuscript writing. All authors had access to the study data and approved the final manuscript.

## DATA AVAILABILITY STATEMENT

The datasets produced in this study have been uploaded to the NCBI’s Sequencing Read Archives (SRA). Health Canada datasets (bam files) are available under BioProject number PRJNA909196 (Reviewer link: https://dataview.ncbi.nlm.nih.gov/object/PRJNA909196?reviewer=aj5t3tmmd0pk60li9pj868mebq). Inotiv-RTP datasets (fastq files) are available under BioProject number PRJNA913202 (Reviewer link: https://dataview.ncbi.nlm.nih.gov/object/PRJNA913202?reviewer=cnjuifkosvjsfqbhk8indv7g02)

## REFERENCES

1. OECD. in OECD Guideline for the Testing of Chemicals, Section 4 (OECD Publishing, Paris, France, 2016).

2. OECD. in Test No. 490: In Vitro Mammalian Cell Gene Mutation Tests Using the Thymidine Kinase Gene 24 (OECD Publishing, Paris, France, 2016).

3. OECD. in OECD Guidelines for the Testing of Chemicals, Section 4 (OECD Publishing, Paris, France, 2020).

4. Salk, J. J. & Kennedy, S. R. Next-generation genotoxicity: Using modern sequencing technologies to assess somatic mutagenesis and cancer risk. Environ Mol Mutagen 61, 135–151 (2020).

5. Beal, M. A. et al. Chemically induced mutations in a MutaMouse reporter gene inform mechanisms underlying human cancer mutational signatures. Commun Biol 3, 438 (2020).

6. Kennedy, S. R. et al. Detecting ultralow-frequency mutations by Duplex Sequencing. Nature Protocols 9, 2586–2606 (2014).

7. Harris, K. L. et al. Quantification of cancer driver mutations in human breast and lung DNA using targeted, error-corrected CarcSeq. Environ Mol Mutagen 61, 872–889 (2020).

8. You, X. et al. Detection of genome-wide low-frequency mutations with Paired-End and Complementary Consensus Sequencing (PECC-Seq) revealed end-repair-derived artifacts as residual errors. Arch Toxicol 94, 3475–3485 (2020).

9. Schmitt, M. W. et al. Detection of ultra-rare mutations by next-generation sequencing. Proc Natl Acad Sci USA 109, 14508–14513 (2012).

10. Abascal, F. et al. Somatic mutation landscapes at single-molecule resolution. Nature 593, 405–410 (2021).

11. Hoang, M. L. et al. Genome-wide quantification of rare somatic mutations in normal human tissues using massively parallel sequencing. Proc Natl Acad Sci USA 113, 9846–9851 (2016).

12. Valentine III, C. C. et al. Direct quantification of in vivo mutagenesis and carcinogenesis using duplex sequencing. Proc Natl Acad Sci USA 117, 52-33414-33425 (2020).

13. LeBlanc, D. P. et al. Duplex sequencing identifies genomic features that determine susceptibility to benzo(a)pyrene-induced in vivo mutations. BMC Genomics 23, 542 (2022).

14. Wang, Y. et al. Genetic toxicity testing using human in vitro organotypic airway cultures: Assessing DNA damage with the CometChip and mutagenesis by Duplex Sequencing. Environ Mol Mutagen 62, 306–318 (2021).

15. Lorge, E. et al. Standardized cell sources and recommendations for good cell culture practices in genotoxicity testing. Mutat Res Fund Mol Mech Mutagen 809, 1–15 (2016).

16. R Core Team. R: A Language and Environment for Statistical Computing. R Foundation for Statistical Computing 4.1.2 (2021).

17. Fox, J. & Weisberg, S. in An R Companion to Applied Regression (Sage, Thousand Oaks, CA, 2019).

18. Hojsgaard, S. & Halekoh, U. doBy: Groupwise Statistics, LSmeans, Linear Contrasts, Utilities. R package version 4.6.11. URL: https://CRAN.R-project.org/package=doBy. (2021).

19. Manders, F. et al. Introduction to MutationalPatterns. URL: https://bioconductor.org/packages/release/bioc/vignettes/MutationalPatterns/inst/doc/Introduction_to_MutationalPatterns.html. (2021).

20. White, P. A., Long, A. S. & Johnson, G. E. Quantitative Interpretation of Genetic Toxicity Dose-Response Data for Risk Assessment and Regulatory Decision-Making: Current Status and Emerging Priorities. Environ Mol Mutatgen 61, 66–83 (2020).

21. Johnson, G. E. in Methods in Molecular Biology (eds Parry, J. & Parry, E.) 55–67 (Springer, New York, NY, 2012).

22. Heflich, R. H. et al. Mutation as a Toxicological Endpoint for Regulatory Decision-Making. Environ Mol Mutagen 61, 34–41 (2020).

23. Saini, N. et al. The Impact of Environmental and Endogenous Damage on Somatic Mutation Load in Human Skin Fibroblasts. PLoS Genet 12, e1006385 (2016).

24. Kucab, J. E. et al. A Compendium of Mutational Signatures of Environmental Agents. Cell 177, 821-836.e16 (2019).

25. Saini, N. et al. UV-exposure, endogenous DNA damage, and DNA replication errors shape the spectra of genome changes in human skin. PLoS Genetics 17, e1009302 (2021).

26. Pleguezuelos-Manzano, C., Puschhof, J., Rosendahl Huber, A. & et al. Mutational signature in colorectal cancer caused by genotoxic pks+ E. coli. Nature 580, 269–273 (2020).

27. Alexandrov, L. & et al. Signatures of mutational processes in human cancer. Nature 500, 415–421 (2013).

28. Alexandrov, L. et al. The repertoire of mutational signatures in human cancer. Nature 578, 94–101 (2020).

29. Kaina, B., Christmann, M., Naumann, S. & Roos, W. P. MGMT: Key node in the battle against genotoxicity, carcinogenicity and apoptosis induced by alkylating agents. DNA Repair 6, 1079–1099 (2007).

30. Nagel, Z. D. et al. Fluorescent reporter assays provide direct, accurate, quantitative measurements of MGMT status in human cells. PLoS One 14, e0208341 (2019).

31. Ehrlich, M., Zhang, X. & Inamdar, N. M. Spontaneous deamination of cytosine and 5-methylcytosine residues in DNA and replacement of 5-methylcytosine residues with cytosine residues. Mutat Res 238, 277–286 (1990).

32. Halliwell, B. Oxidative stress in cell culture: an under-appreciated problem? FEBS Lett 540, 3–6 (2003).

33. Markkanen, E. Not breathing is not an option: How to deal with oxidative DNA damage. DNA Repair 59, 82–105 (2017).

