## Supplementary Figure S6 for "Error-corrected Duplex Sequencing enables direct detection and quantification of mutations in human TK6 cells with remarkable inter-laboratory consistency"

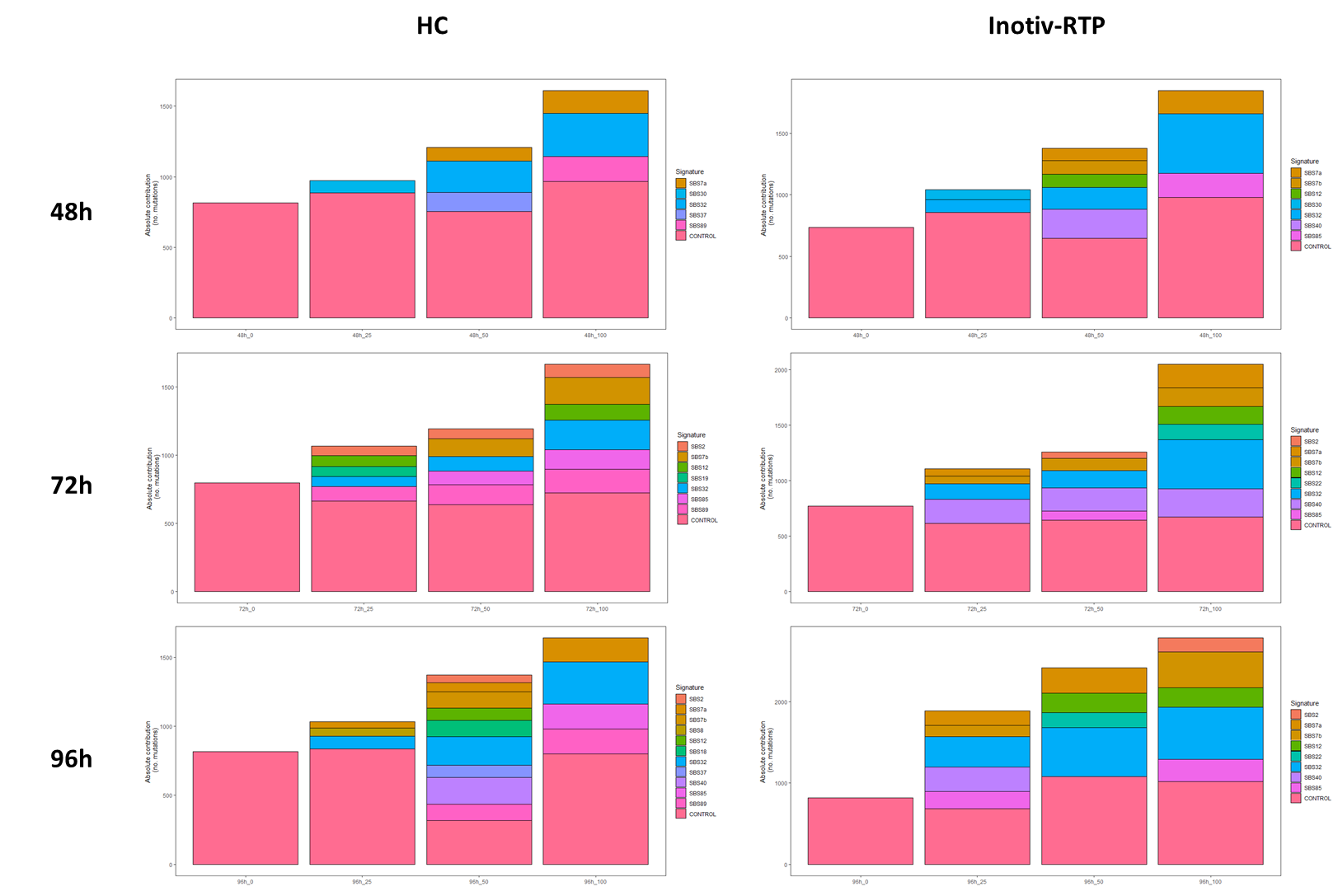

Supplementary Figure S6. Contribution of mutations to the mutational signatures constituting the reconstructed trinucleotide spectra of TK6 cells exposed to ENU at HC and Inotiv-RTP. Control mutational signatures (pink) were generated using the trinucleotide spectra in the vehicle controls at each time point. The stacked bars represent the combination of signatures from the Catalogue of Somatic Mutations in Cancer (COSMIC) with the highest cosine similarity to the original trinucleotide spectra in TK6 cells at each concentration and time point.
