## Supplementary Table S1 for "Error-corrected Duplex Sequencing enables direct detection and quantification of mutations in human TK6 cells with remarkable inter-laboratory consistency"

Supplementary Table S1. TwinStrand Human mutagenesis panel target characteristics

| **Target** | **Start Location** | **End Location** | **Location Relative to Genes** | **Gene (NCBI Genome Data Viewer)** |
| --- | --- | --- | --- | --- |
| chr11 | 108510787 | 108513187 | Genic | *EXPH5* |
| chr13 | 75803912 | 75806312 | Genic | *LMO7* |
| chr14 | 74661755 | 74664155 | Genic | *AREL1* |
| chr18 | 5749264 | 5751664 | Genic | *MIR3976HG* |
| chr2 | 40162767 | 40165167 | Genic | *SLC8A1-AS1* |
| chr20 | 24153684 | 24156084 | Genic | *LINC01721* |
| chr6 | 155239014 | 155241414 | Genic | *TIAM2* |
| chr7 | 11732774 | 11735174 | Genic | *THSD7A* |
| chr8 | 51513056 | 51515456 | Genic | *PXDNL* |
| chr9 | 23709463 | 23711863 | Genic | *ELAVL2* |
| chr1 | 84597127 | 84599527 | Genic/ intergenic | *LINC01461* |
| chr16 | 51754103 | 51756503 | Genic/ intergenic | *LOC105371257* |
| chr19 | 31831021 | 31833421 | Genic/ intergenic | *LINC01837* |
| chr4 | 22386244 | 22388644 | Genic/ intergenic | *ADGRA3* |
| chr12 | 114115043 | 114117443 | Intergenic |  |
| chr15 | 46089737 | 46092137 | Intergenic |  |
| chr17 | 70672726 | 70675126 | Intergenic |  |
| chr21 | 23665976 | 23668376 | Intergenic |  |
| chr22 | 48262370 | 48264770 | Intergenic |  |
| chr10 | 128969037 | 128971437 | Intergenic |  |
