## Supplementary Table S2 for "Error-corrected Duplex Sequencing enables direct detection and quantification of mutations in human TK6 cells with remarkable inter-laboratory consistency"

Supplementary Table S2. Mutation frequency at each concentration and time point (n=2) in TK6 cells exposed to ENU measured by Duplex Sequencing.

|  | **HC** | | | | **Inotiv-RTP** | | | |
| --- | --- | --- | --- | --- | --- | --- | --- | --- |
|  | 0 µM | 25 µM | 50 µM | 100 µM | 0 µM | 25 µM | 50 µM | 100 µM |
| **48h** | 5.37 x 10^-7^ | 7.44 x 10^-7^ | 8.85 x 10^-7^ | 9.76 x 10^-7^ | 4.58 x 10^-7^ | 6.17 x 10^-7^ | 7.96 x 10^-7^ | 10.5 x 10^-7^ |
|  | 5.64 x 10^-7^ | 7.08 x 10^-7^ | 8.45 x 10^-7^ | 10.9 x 10^-7^ | 4.77 x 10^-7^ | 6.39 x 10^-7^ | 7.37 x 10^-7^ | 11.2 x 10^-7^ |
| **72h** | 5.98 x 10^-7^ | 6.75 x 10^-7^ | 8.87 x 10^-7^ | 10.5 x 10^-7^ | 4.77 x 10^-7^ | 6.37 x 10^-7^ | 8.01 x 10^-7^ | 11.2 x 10^-7^ |
|  | 5.29 x 10^-7^ | 7.26 x 10^-7^ | 9.14 x 10^-7^ | 11.3 x 10^-7^ | 4.57 x 10^-7^ | 6.77 x 10^-7^ | 7.89 x 10^-7^ | 11.6 x 10^-7^ |
| **96h** | 5.48 x 10^-7^ | 6.51 x 10^-7^ | 8.51 x 10^-7^ | 10.1 x 10^-7^ | 4.8 x 10^-7^ | 5.97 x 10^-7^ | 8.08 x 10^-7^ | 12.3 x 10^-7^ |
|  | 5.54 x 10^-7^ | 7.18 x 10^-7^ | 9.29 x 10^-7^ | 10.8 x 10^-7^ | 4.45 x 10^-7^ | 6.35 x 10^-7^ | 8.21 x 10^-7^ | 11.9 x 10^-7^ |
