## Supplementary Table S3 for "Error-corrected Duplex Sequencing enables direct detection and quantification of mutations in human TK6 cells with remarkable inter-laboratory consistency"

Supplementary Table S3. Summary of the Analysis of Deviance Table of the HC and Inotiv-RTP datasets.

| Parameter | LR Chi-squared value | Df | Pr(>Chisq) | Statistical Significance |
| --- | --- | --- | --- | --- |
| Concentration | 1880.48 | 3 | < 2.2E-16 | *** |
| Laboratory | 15.96 | 1 | 0.00006486 | *** |
| Time Point | 4.37 | 2 | 0.1124 |  |
| Concentration : Laboratory | 60.17 | 3 | 5.407E-13 | *** |
| Concentration : Time Point | 6.66 | 6 | 0.3535 |  |
| Laboratory : Time Point | 2.75 | 2 | 0.2523 |  |
| Concentration : Laboratory : Time Point | 3.82 | 6 | 0.7012 |  |

*** p-value < 0.001 (Holm-Sidak adjusted)

The results of the Analysis of Deviance Table for the overall MF data from HC and Inotiv-RTP are summarized in Supplementary Table S2. In the pooled analysis, the effects of ENU concentration and laboratory (HC vs. Inotiv-RTP) on MF, as well as the interaction between concentration and laboratory were significant in the model. The Likelihood Ratio (LR) test statistic indicated highly significant and prominent concentration-related effects on MF. There was no evidence of a time point-related effect on MF or interactions of other parameters with time point.
