## Supplementary Table S4 for "Error-corrected Duplex Sequencing enables direct detection and quantification of mutations in human TK6 cells with remarkable inter-laboratory consistency"

Supplementary Table S4. The Bonferroni-corrected results of the Analysis of Deviance Table on individual target sites of the human mutagenesis panel.

|  | Chromosome | | | | | | | | | | | | | | | | | | | | | | | | | | | | | | | | | | | |
| --- | --- | --- | --- | --- | --- | --- | --- | --- | --- | --- | --- | --- | --- | --- | --- | --- | --- | --- | --- | --- | --- | --- | --- | --- | --- | --- | --- | --- | --- | --- | --- | --- | --- | --- | --- | --- |
|  | 1 | 2 | 4 | | 6 | | 7 | | 8 | | 9 | | 10 | | 11 | | 12 | | 13 | | 14 | | 15 | | 16 | 17 | | 18 | | | 19 | 20 | | 21 | | 22 |
| Concentration | *** | *** | *** | | *** | | *** | | *** | | *** | | *** | | *** | | *** | | *** | | *** | | *** | | *** | *** | | *** | | | *** | *** | | *** | | *** |
| Laboratory |  |  | |  | |  | |  | |  | |  | |  | |  | |  | |  | |  | |  | | | *** | |  |  | | | *** | | *** |  |
| Time Point |  | ** |  | |  | |  | |  | |  | |  | |  | |  | |  | |  | |  | |  |  | |  | | |  |  | |  | |  |
| Concentration : Laboratory |  |  | *** | |  | |  | |  | |  | |  | | ** | |  | |  | |  | |  | |  |  | |  | | |  |  | |  | |  |

The HC and Inotiv-RTP datasets were pooled for this analysis.

* p-value 0.01-0.05

** p-value 0.001-0.01

*** p-value 0-0.001

The target sites were then analyzed individually to study the observed significant effects in more detail and determine whether there were consistencies across the 20 sites that support the effects of concentration and laboratory, and the concentration-laboratory interactions observed in the pooled analysis. Significant concentration-related effects were observed at all 20 target sites, consistent with the overall results (Supplementary Table S3). However, only three target sites on chromosomes 17, 20, and 21 showed significant laboratory-related effects on MF, while at only two sites on chromosomes 4 and 11, an interaction between laboratory and concentration occurred at the highest exposure concentration. Only the target on chromosome 2 was affected by time point. Consistent with the pooled analysis, no interactions of time point with any other parameters were identified for any of the target sites.
