## Supplementary Table S5 for "Error-corrected Duplex Sequencing enables direct detection and quantification of mutations in human TK6 cells with remarkable inter-laboratory consistency"

Supplementary Table S5. COSMIC signatures that showed concentration-dependent increases in cosine similarity in TK6 cells exposed to ENU

|  |  | **SBS7a** | **SBS7b** | **SBS11** | **SBS30** | **SBS32** |
| --- | --- | --- | --- | --- | --- | --- |
| HC | 48h_0 | 0.45 | 0.51 | 0.41 | 0.50 | 0.46 |
|  | 48h_25 | 0.48 | 0.53 | 0.48 | 0.57 | 0.53 |
|  | 48h_50 | 0.53 | 0.59 | 0.56 | 0.63 | 0.64 |
|  | 48h_100 | 0.60 | 0.66 | 0.62 | 0.70 | 0.67 |
|  | 72h_0 | 0.42 | 0.54 | 0.52 | 0.55 | 0.53 |
|  | 72h_25 | 0.50 | 0.55 | 0.50 | 0.60 | 0.58 |
|  | 72h_50 | 0.55 | 0.62 | 0.55 | 0.63 | 0.58 |
|  | 72h_100 | 0.57 | 0.65 | 0.60 | 0.65 | 0.63 |
|  | 96h_0 | 0.41 | 0.49 | 0.39 | 0.52 | 0.47 |
|  | 96h_25 | 0.47 | 0.54 | 0.48 | 0.57 | 0.55 |
|  | 96h_50 | 0.58 | 0.62 | 0.59 | 0.67 | 0.63 |
|  | 96h_100 | 0.56 | 0.61 | 0.59 | 0.66 | 0.66 |
| Inotiv-RTP | 48h_0 | 0.45 | 0.51 | 0.40 | 0.48 | 0.43 |
|  | 48h_25 | 0.49 | 0.57 | 0.50 | 0.57 | 0.56 |
|  | 48h_50 | 0.58 | 0.64 | 0.55 | 0.62 | 0.57 |
|  | 48h_100 | 0.53 | 0.58 | 0.60 | 0.65 | 0.68 |
|  | 72h_0 | 0.43 | 0.47 | 0.38 | 0.48 | 0.46 |
|  | 72h_25 | 0.56 | 0.61 | 0.57 | 0.63 | 0.62 |
|  | 72h_50 | 0.54 | 0.57 | 0.54 | 0.62 | 0.61 |
|  | 72h_100 | 0.61 | 0.66 | 0.65 | 0.69 | 0.69 |
|  | 96h_0 | 0.48 | 0.55 | 0.53 | 0.61 | 0.58 |
|  | 96h_25 | 0.51 | 0.57 | 0.47 | 0.55 | 0.51 |
|  | 96h_50 | 0.56 | 0.61 | 0.58 | 0.67 | 0.67 |
|  | 96h_100 | 0.62 | 0.67 | 0.66 | 0.70 | 0.69 |

SBS7a,b - UV exposure

SBS11	- Temozolomide (alkylator) exposure

SBS30	- Defective BER due to NTHL1 (endonuclease III) deficiency

SBS32	- Immunosuppression via aziothioprine exposure
