## Supplementary Discussion for "Error-corrected Duplex Sequencing enables direct detection and quantification of mutations in human TK6 cells with remarkable inter-laboratory consistency"

We investigated what additional mechanistic insight we could gain from the mutation spectra data (Supplementary Table S5 and Supplementary Figure S6). Two COSMIC signatures with proposed etiologies that contributed to the ENU-induced TK6 trinucleotide mutation spectra were SBS7a and b and SBS32, which are associated with UV exposure and immunosuppression induced by azathioprine treatment, respectively ^1, 2^. While both SBS7b and SBS32 were present in both untreated and treated TK6 cells, the contribution of both signatures to the mutation spectra increased with ENU concentration (Supplementary Figure S6). UV exposure induces C:G > T:A mutations, which are also induced by ENU although through different mutagenic adducts^3^. Azathioprine is a metabolic precursor to 6-thiopurine, a nucleoside analogue that can be incorporated into DNA and induce clastogenic and mutagenic effects^4^. In the analysis of cosine similarity between COSMIC signatures and the TK6 mutation spectra, an additional two signatures, SBS11 and SBS30, were observed to increase in similarity as the concentration of ENU increased. SBS11 is associated with exposure to temozolomide, another alkylating agent. Interestingly, SBS30 may indicate defective base excision repair due to a deficiency in NTH1, a glycosylase that initiates the repair of mainly oxidized pyrimidines^5, 6^. There is insufficient information on the level of NTH1 activity in TK6 cells and, thus, it is not possible to confirm whether the cosine similarity in TK6 spectra to this mutational signature aligns with direct observations in the cell line^7^. Nonetheless, the current results are a promising indication that mutational signatures found in human cancer genomes can be detected in TK6 cells and, thus, support the findings in previous studies that insights into chemical carcinogenicity could potentially be gained through *in vitro* testing^8^.

**References**

1. Hayward, N. *et al*. Whole-genome landscapes of major melanoma subtypes. *Nature* **54**, 175-180 (2017).

2. Inman, G. J. *et al*. The genomic landscape of cutaneous SCC reveals drivers and a novel azathioprine associated mutational signature. *Nat Commun* **9**, 3667 (2018).

3. Ikehata, H. & Ono, T. The Mechanisms of UV Mutagenesis. *J Radiat Res* **52**, 115-125 (2011).

4. Melo-Bisneto, A. V. *et al*. Recombinogenic, genotoxic, and cytotoxic effects of azathioprine using in vivo assays. *J Toxicol Environ Health* **84**, 261-271 (2021).

5. Nik-Zainal, S. *et al*. Mutational processes molding the genomes of 21 breast cancers. *Cell* **149**, 979-993 (2012).

6. Alexandrov, L. & et al. Signatures of mutational processes in human cancer. *Nature* **500**, 415-421 (2013).

7. Yang, N., Chaudhry, A. & Wallace, S. Base excision repair by hNTH1 and hOGG1: A two edged sword in the processing of DNA damage in γ-irradiated human cells. *DNA Repair* **5**, 43-51 (2006).

8. Kucab, J. E. *et al*. A Compendium of Mutational Signatures of Environmental Agents. *Cell* **177**, 821-836.e16 (2019).
