## Supplementary Figure S1 for "Error-corrected Duplex Sequencing enables direct detection and quantification of mutations in human TK6 cells with remarkable inter-laboratory consistency"

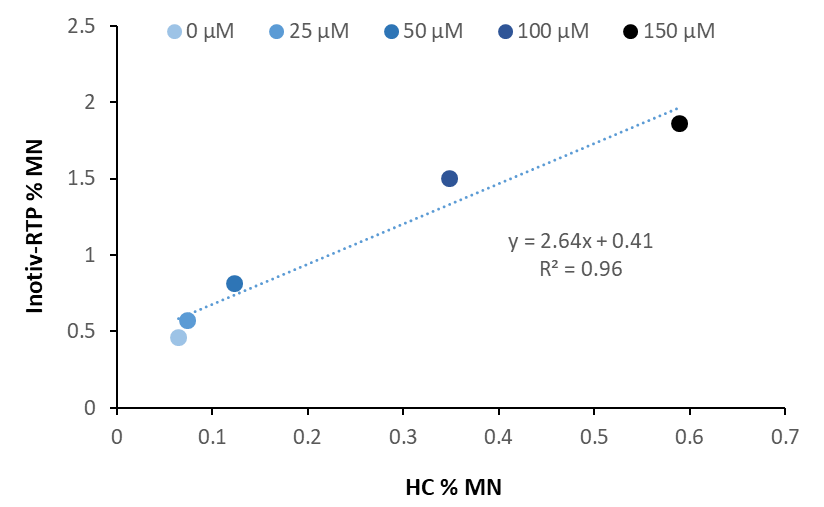

Supplementary Figure S1. Linear correlation of the micronucleus frequency in TK6 cells after a 24h exposure to ENU, measured at HC (n=4) and Inotiv-RTP (n=2).
