## Supplementary Figure S2 for "Error-corrected Duplex Sequencing enables direct detection and quantification of mutations in human TK6 cells with remarkable inter-laboratory consistency"

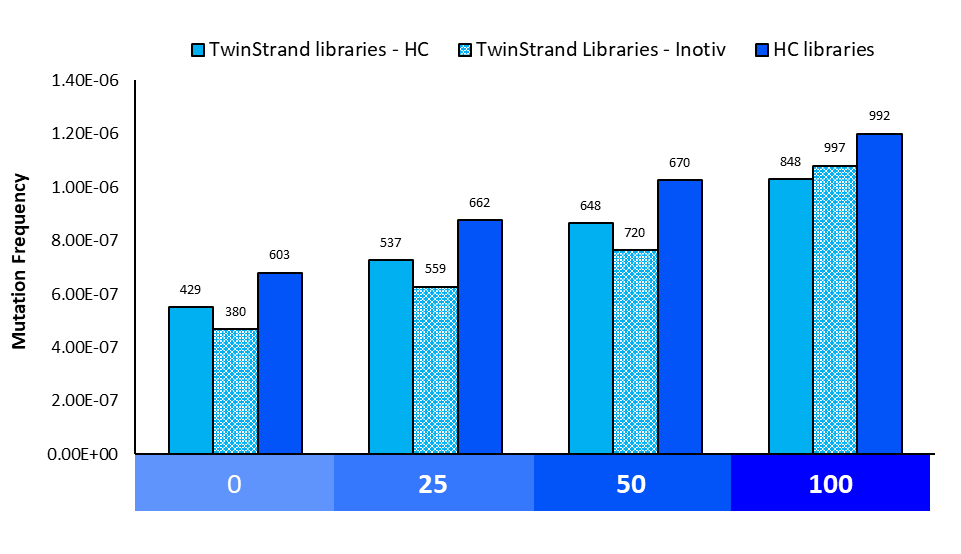

Supplementary Figure S2. Mutation frequency in TK6 cells exposed to ENU measured by Duplex Sequencing (TwinStrand Biosciences). Two replicate sets of 48 h DNA samples were retained for in-house DuplexSeq library preparation at HC (n=2).
