## Supplementary Figure S3 for "Error-corrected Duplex Sequencing enables direct detection and quantification of mutations in human TK6 cells with remarkable inter-laboratory consistency"

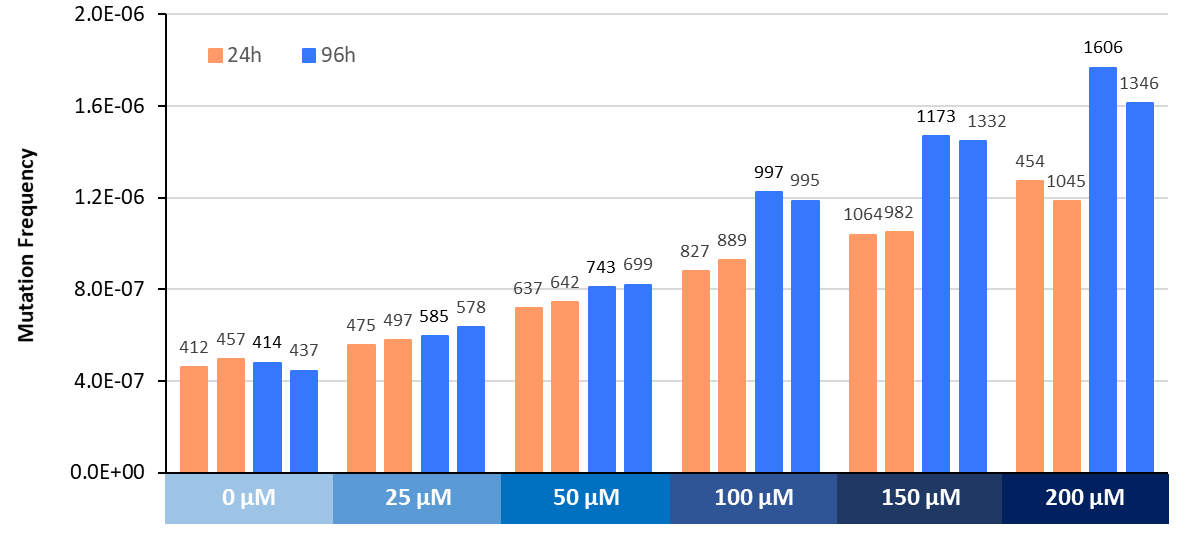

Supplementary Figure 3. Mutation frequency in TK6 cells exposed to ENU at Inotiv-RTP measured by Duplex Sequencing (TwinStrand Biosciences). Cells were sampled 24 and 96 hours following the initial exposure (n=2). These samples were excluded from the comparison with HC results because a 24h sampling time and ENU concentrations of 150 µM and 200 µM were not part of the HC study.
