## Supplementary Figure S4 for "Error-corrected Duplex Sequencing enables direct detection and quantification of mutations in human TK6 cells with remarkable inter-laboratory consistency"

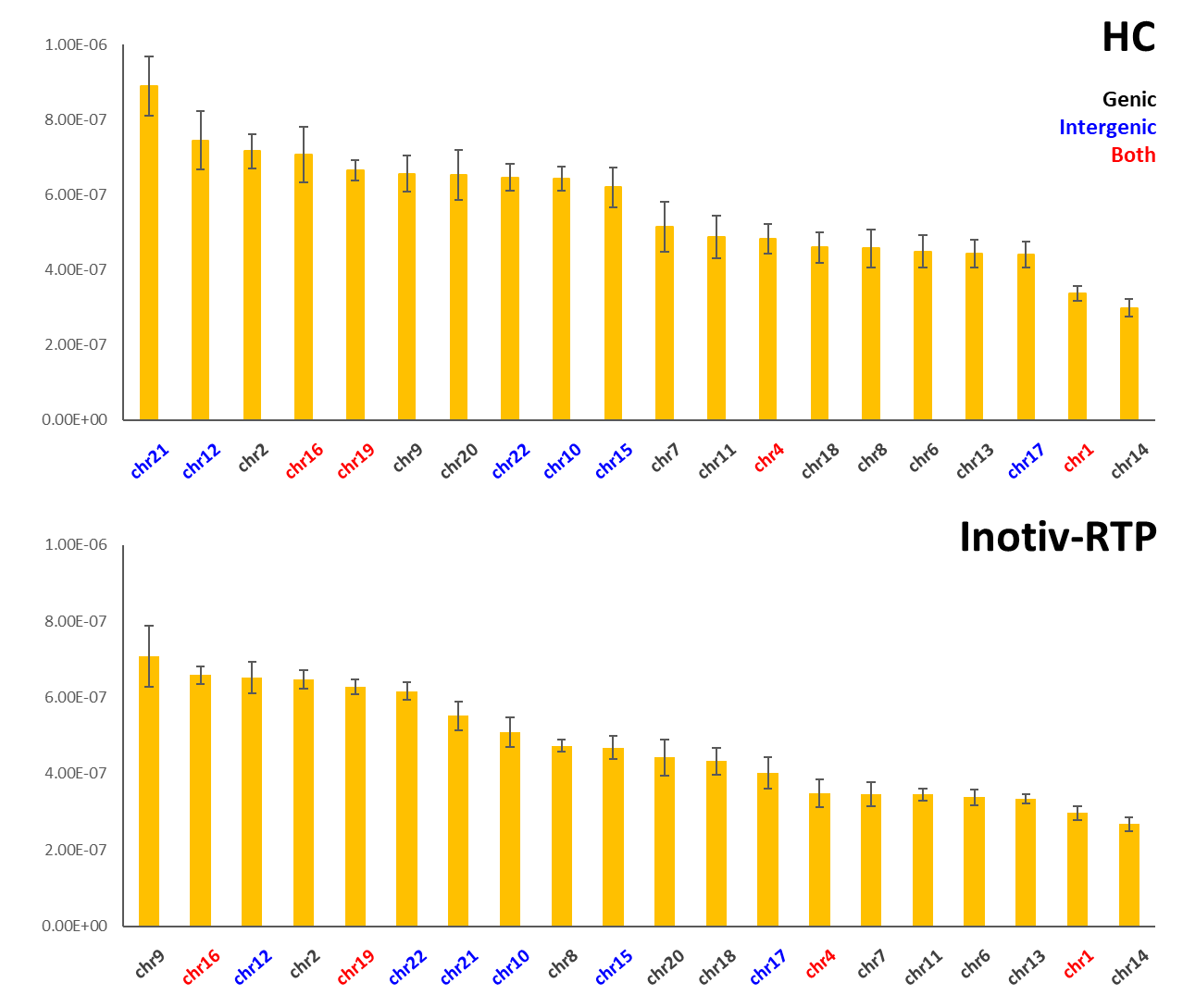

Supplementary Figure S4. Mutation frequency in the 20 target sites of the Duplex Sequencing human mutagenicity panel measured in DMSO-treated TK6 cells (vehicle control) at HC and Inotiv-RTP. The mutation frequency at 48, 72, and 96h were averaged (n=2 at each time point; n=6 in total). Black indicates genic sites, blue indicates intergenic sites, and red indicates those that include both genic and intergenic sequences. The target sites are ordered by mutation frequency, from the highest (left) to the lowest (right). Error bars represent standard error.
