## Supplementary Figure S5 for "Error-corrected Duplex Sequencing enables direct detection and quantification of mutations in human TK6 cells with remarkable inter-laboratory consistency"

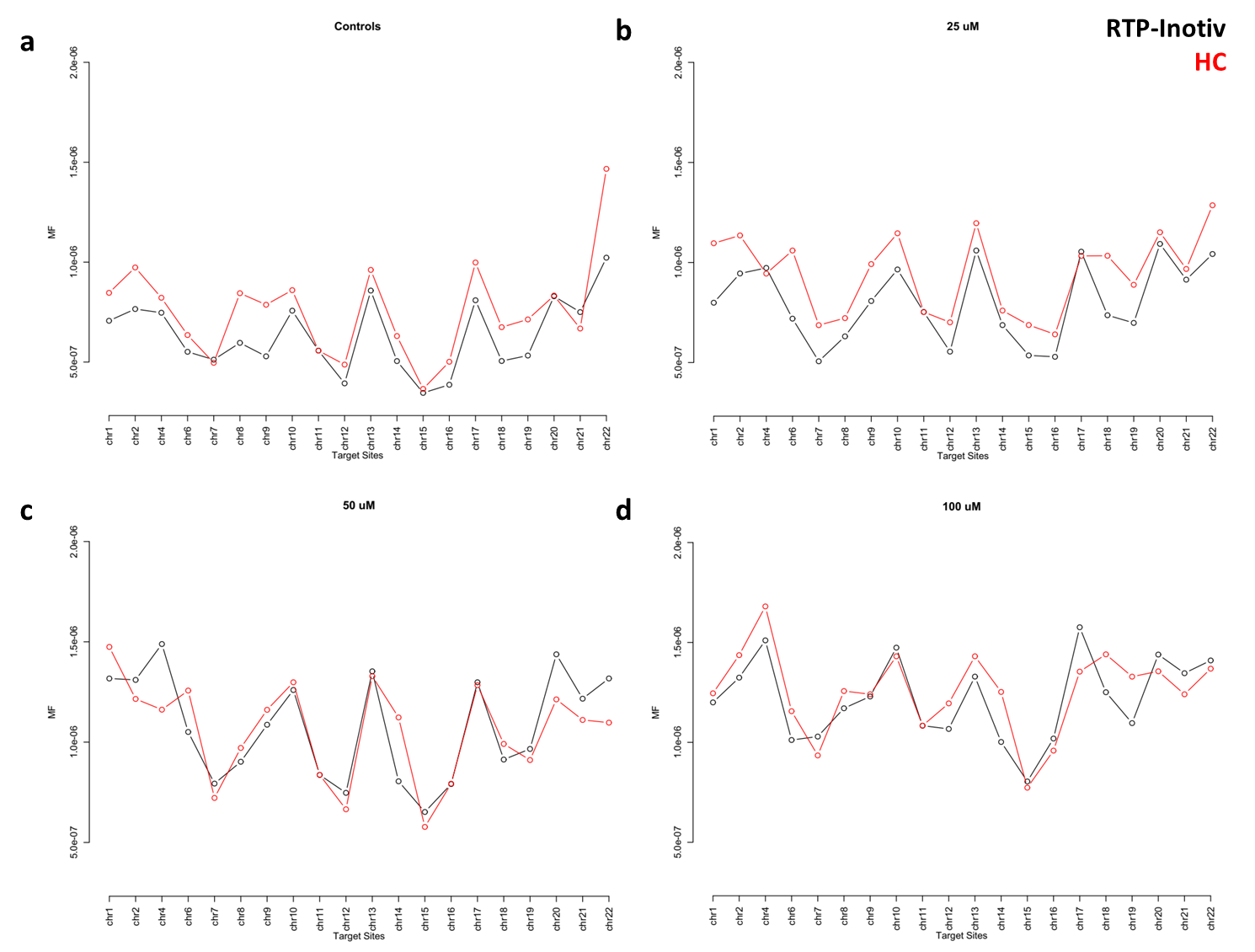

Supplementary Figure S5. Factor plots of the generalized linear mixed models (GLMM) of mutation frequency across the 20 target sites of the Duplex Sequencing human mutagenicity panel at different concentrations of ENU: 0 (a), 25 (b), 50 (c), and 100 µM (d). Black represents Inotiv-RTP data and red represents HC data. The Holm-Sidak adjustment has been applied to correct for multiple comparisons.
